# Comparison between diffusion MRI tractography and histological tract-tracing of cortico-cortical structural connectivity in the ferret brain

**DOI:** 10.1101/517136

**Authors:** C. Delettre, A. Messé, L-A. Dell, O. Foubet, K. Heuer, B. Larrat, S. Meriaux, J-F. Mangin, I. Reillo, C. de Juan Romero, V. Borrell, R. Toro, C. C. Hilgetag

## Abstract

The anatomical wiring of the brain is a central focus in network neuroscience. Diffusion MRI tractography offers the unique opportunity to investigate the brain fiber architecture in vivo and non invasively. However, its reliability is still highly debated. Here, we explored the ability of diffusion MRI tractography to match invasive anatomical tract-tracing connectivity data of the ferret brain. We also investigated the influence of several state-of-the-art tractography algorithms on this match to ground truth connectivity data. Tract-tracing connectivity data were obtained from retrograde tracer injections into the occipital, parietal and temporal cortices of adult ferrets. We found that the relative densities of projections identified from the anatomical experiments were highly correlated with the estimates from all the studied diffusion tractography algorithms (Spearman’s rho ranging from 0.67 to 0.91), while only small, non-significant variations appeared across the tractography algorithms. These results are comparable to findings reported in mouse and monkey, increasing the confidence in diffusion MRI tractography results. Moreover, our results provide insights into the variations of sensitivity and specificity of the tractography algorithms and hence, into the influence of choosing one algorithm over another.

## Introduction

Brain function emerges from the communication of spatially distributed large-scale networks via the underlying structural connectivity architecture (Engel, Gerloff, Hilgetag, & Nolte, 2013; Kandel, Schwartz, Jessell, Siegelbaum, & Hudspeth, 2012; Park & Friston, 2013; Varela, Lachaux, Rodriguez, & Martinerie, 2001). Systematic analysis of structural connectivity has revealed characteristic features of brain networks, including the presence of modules, hubs and higher-order topological properties, thought to support efficient information processing (Sporns, 2010). Moreover, structural connectivity is considered as a neural substrate that is affected in various pathological conditions, such as Alzheimer’s disease and schizophrenia spectrum disorders (Fornito & Bullmore, 2015). Therefore, reliable estimates of brain structural connectivity are essential for advancing our understanding of the network basis of brain function.

Diffusion MRI tractography is an indirect approach for inferring brain structural connectivity from the brownian motion of water molecules constrained by the axonal fiber architecture (Jeurissen, Descoteaux, Mori, & Leemans, 2017). Thus, it provides the unique opportunity to investigate, *in vivo* and non invasively, the structural connectivity of intact or altered brains, such as in the case of stroke (Visser et al., 2018), in longitudinal analysis of brain development (Hagmann et al., 2010) or *in utero* acquisitions of prenatal brain structure (Kasprian et al., 2008). However, the reliability of diffusion MRI tractography for properly mapping structural connections remains highly debated (Jones, Knösche, & Turner, 2013; Thomas et al., 2014). Therefore, validation appears as a key step in evaluating current methodologies and identifying new perspectives of improvement (Dyrby, Innocenti, Bech, & Lundell, 2018).

A small number of studies designed benchmarks in order to explore the reliability of diffusion MRI tractography (Schilling et al., 2018). For example, using a phantom dataset composed of known tracts reconstructed by diffusion MRI tractography as ground truth, the accuracy of a large number of state-of-the-art tractography algorithms was assessed in humans (Maier-Hein et al., 2017). The results showed, for all the algorithms, their ability to recover most of the existing bundles, but also revealed a variable, but substantial, number of false positives. Similarly (Sarwar, Ramamohanarao, & Zalesky, 2018) compared deterministic and probabilistic tractography algorithms with a numerically generated phantom and concluded on a trade-off to be made between sensitivity and specificity depending on the type of tractography algorithm. While these studies provided a first estimate of the specificity and sensitivity of a wide range of tractography algorithms, the ground truths used were based on diffusion MRI tractography or numerically generated and thus one can debate their realism.

To date, the gold standard for assessing structural brain connectivity is provided by tract-tracing experiments, which physically investigate, at the cellular level, the relative number of connections of an area to the rest of the brain using viral, bacterial or biotinylated dextran agents (Bizley, Bajo, Nodal, & King, 2015; Bota, Sporns, & Swanson, 2015; Markov et al., 2014; Zingg et al., 2015). These agents act as either anterograde or retrograde tracers. Such histological tracing of anatomical connections provides directional (as well as laminar) information on projection. In the case of retrograde tracing, histological tracing also quantifies the number of axons in a projection, since each labeled projection neuron provides one axon. Studies performed in macaque (Azadbakht et al., 2015; Donahue et al., 2016; Schilling, Nath, et al., 2019; Zhang et al., 2018), squirrel monkey (Gao et al., 2013; Schilling, Gao, et al., 2019), pig (Knösche, Anwander, Liptrot, & Dyrby, 2015), mouse (Calabrese, Badea, Cofer, Qi, & Johnson, 2015) and rat (Sinke et al., 2018), have explored the relationship between tract-tracing experiments and tractography. Overall these studies have shown that diffusion MRI tractography provides a good estimate of structural brain connectivity. Few explorations have been made on the ability of the different tractography approaches available to estimate structural connectivity weights (Gao et al., 2013). Previous studies have mainly focused on the ability of tractography algorithms to properly estimate white-matter pathways by means of voxel-wise overlap (Knösche et al., 2015), or on the detectability (presence or absence) of connections (Sinke et al., 2018), or both (Schilling, Gao, et al., 2019; Schilling, Nath, et al., 2019).

The ferret brain starts to fold after birth, and reaches the adult folding pattern at about one month of age. Therefore, the ferret is a widely used animal model for studying brain folding (Barnette et al., 2009; Feng, Clayton, Chang, Okamoto, & Bayly, 2013). Furthermore, these mammals display complex behavior, are inexpensive to house and have a short gestation period as well as a limited life time, making them an attractive ‘whole lifespan model’ (Fox, 1998). Recently established extensive tract-tracing connectivity data of the ferret (Dell, Innocenti, Hilgetag, & Manger, 2019a, 2019b, 2019c) have made it possible to compare anatomical cortical connectivity with that reported by tractography methods. Thus, the present study aimed to use the ferret as an animal model to assess the performance of six diffusion tractography algorithms compared to histological tract-tracing data from the occipital, parietal and temporal cortices in the ferret. Overall, our results showed that diffusion MRI tractography provides statistically significant estimates of ferret brain structural connectivity, although the different tractography algorithms presented variations in terms of sensitivity and specificity.

## Material and Methods

### Ferret brain atlas

We used a parcellation based on the atlas of the posterior cortex by (Bizley & King, 2009). The parcellation scheme was manually drawn on the left hemisphere in the diffusion MRI space using the online tool BrainBox (Heuer, Ghosh, Sterling, & Toro, 2016), http://brainbox.pasteur.fr/). Tract-tracing data were available for areas 17, 18, 19, 21 (occipital visual areas); 20a and 20b combined (temporal visual areas); PPr and PPc (parietal visual areas) (Figure 1A).

**Figure 1:**
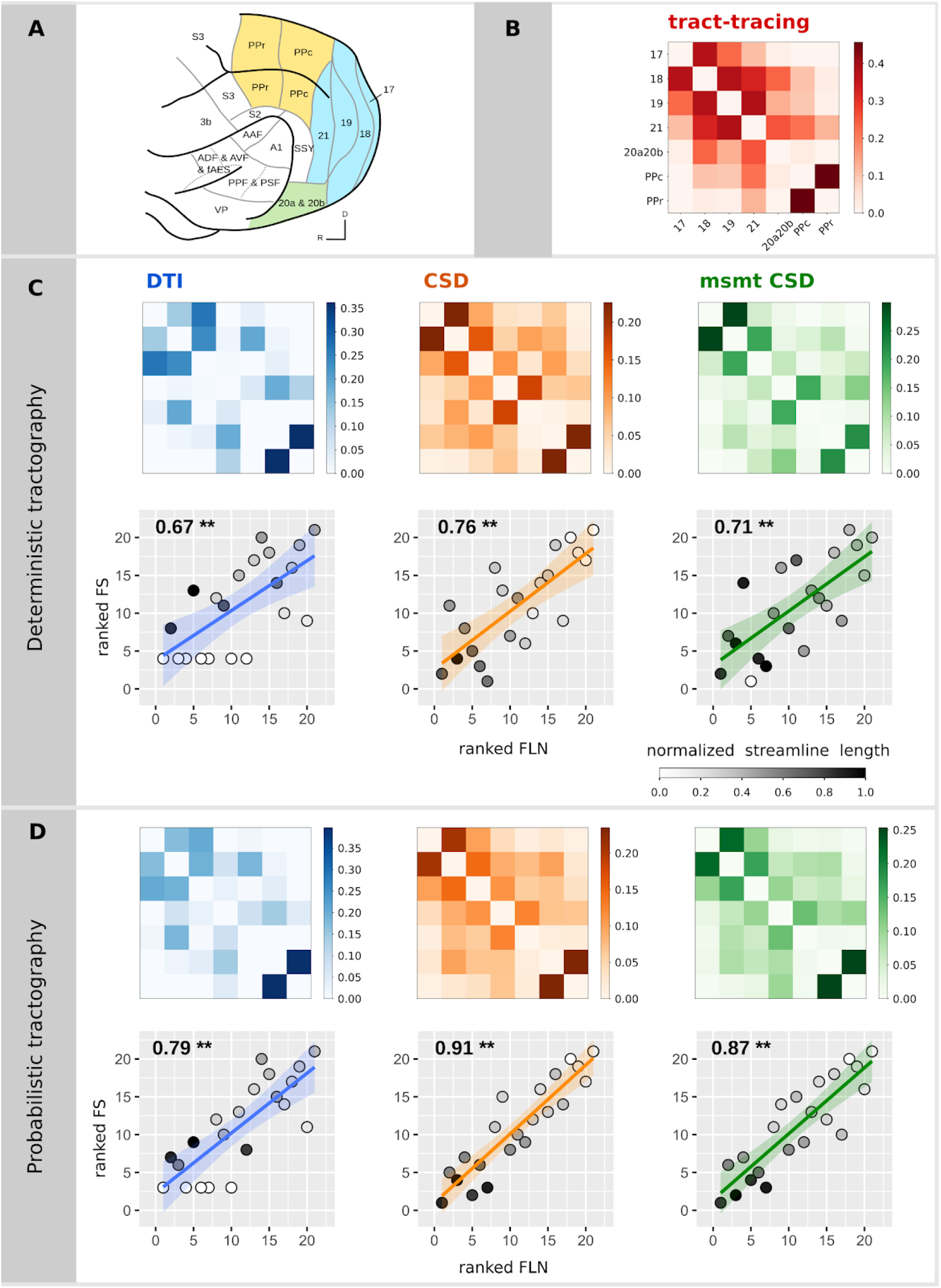
Relationship between diffusion MRI tractography and tract-tracing experiments. (A) Ferret brain atlas according to the parcellation of Bizley and King (Figure adapted from (Bizley & King, 2009)). The regions of interest for the comparative study are those colored. Colors code for the different visual brain areas: posterior parietal (yellow), occipital (blue) and temporal cortices (green). (B) Structural connectivity matrix based on tract-tracing experiments, where the weights represent the fraction of labeled neurons (FLN). Structural connectivity matrices estimated from the deterministic (C) and the probabilistic (D) tractography algorithms and the associated scatterplots of the ranked FLN vs. the ranked FS. Each point in the scatter plot corresponds to a connection between a pair of areas for the tract-tracing results (abscissa) and the diffusion results (ordinate). The ranked weights of the connections allow to visualize of the Spearman’s rho as the slope of the fitted curve. Grey colors code for the average streamline length (values normalized by the maximum streamline length of all the algorithms). P-values inferior to 1.10^−3^ are indicated by **.

### Diffusion MRI data

High resolution MRI were acquired *ex vivo* using a small animal 7 Tesla Bruker MRI scanner (Neurospin, Saclay, France). The acquisitions were performed *post mortem* in order to improve sensitivity (Holmes et al., 2017). The brain was obtained from a 2 month old ferret. The ferret was euthanized by an overdose of pentobarbital and perfused transcardially with 0.9% saline solution and post-fixed with phosphate-buffered 4% paraformaldehyde (PFA). After extraction, the brain was stored at 4°C in a 4% PFA solution until the MRI acquisition. All procedures were approved by the IACUC of the Universidad Miguel Hernández and CSIC, Alicante, Spain.

24 hours before MRI acquisition, the brain was transferred to a 0.01M Phosphate Buffer Saline (PBS) solution for rehydration. Shortly before MRI acquisition, the brain sample was transferred to a plastic tube filled with non protonic liquid (fluorinert) in order to avoid air-tissue interfaces that may induce susceptibility artefacts, as well as to avoid foldover MRI artefacts due to proton signal coming from a protonic liquid outside the imaging field of view (McRobbie, Moore, & Graves, 2017). The tube was then placed in a dedicated holder in the middle of the transmit/receive MRI volume radiofrequency coil. Temperature stability was ensured by a regulated room temperature as well as the cooling of the gradient coils, by water at 16°C was constantly flowing inside the innermost part of the magnet. The equilibrium temperature at the sample was of 20°C.

High-resolution T2-weighted MRI data was acquired using a multi-slice multi echo (MSME) sequence with 18 echo times and 0.12 mm isotropic voxels. Diffusion MRI data was acquired using a multislice 2D spin-echo segmented DTI-EPI sequence (4 segments) and the following parameters: TR = 40 s; TE = 32 ms; matrix size = 160 × 120 × 80; 0.24 mm isotropic voxels; 200 diffusion-weighted directions with b = 4000 s/mm^2^; and 10 b0 at the beginning of the sequence, diffusion gradient duration = 5 ms and diffusion gradient separation = 17 ms. Thanks to strong gradients compared to clinical scanners, the b-value could reach 4000 s/mm^2^ ensuring a strong diffusion weighting and therefore an improved sensitivity to anisotropy while keeping the echo time low enough to save SNR and limit EPI distortions. A b-value of 4000 s/mm^2^ has been previously shown to be a good compromise for disentangling crossing fibers for ex-vivo imaging (Dyrby et al., 2011). The noise introduced by the high diffusion weighting was balanced by a high angular resolution. The 200 directions were generated as non collinear directions uniformly distributed over a sphere (Hasan, Parker, & Alexander, 2001). The spatial resolution was chosen as the highest resolution available on the scanner in order to achieve a good SNR while keeping a reasonable acquisition time. We obtained an SNR of 4.2, measured as the ratio between the mean of our signal in the brain and the standard deviation of the signal in the background. The total acquisition time of the diffusion MRI sequences was about 37 hours.

### Preprocessing

First, MRI data were converted from the 2dseq bruker format to the standard NIFTI format using a modified version of the bruker2nifti script (original version: https://github.com/SebastianoF/bruker2nifti; modified version: https://github.com/neuroanatomy/bruker2nifti). For a limited number of volumes, EPI trajectories were poorly corrected, by the Bruker routine image reconstructor, which resulted in noisy volumes. In order to exclude these volumes, diffusion-weighted directions for which their mean signal was two standard deviations away from the global average across all the volumes were visually inspected and removed. Three out of 200 volumes were removed following this criterion. The preprocessing steps were mainly done using MRtrix3 functions and included: a local principal component analysis (LPCA) denoising (Veraart et al., 2016), Gibbs ringing correction (Kellner, Dhital, Kiselev, & Reisert, 2016), FSL-based eddy current correction (Andersson & Sotiropoulos, 2016; Jenkinson, Beckmann, Behrens, Woolrich, & Smith, 2012) and B1 field inhomogeneity correction (Tustison et al., 2010). A brain mask was manually segmented from the high-resolution T2 image, in order to obtain a precise delineation of the sulci and gyri. Spatial normalization using a linear transformation between the high-resolution T2 volume and diffusion MRI data was computed using FLIRT tools (Jenkinson, Bannister, Brady, & Smith, 2002), and the brain mask was registered to the diffusion space.

### Tractography

We evaluated the ability of different tractography approaches to reliably reconstruct structural connectivity provided by the tract-tracing experiments. We considered three local models: (1) the diffusion tensor (DT) model; (2) fiber orientation distribution (FOD) estimated with a constrained spherical deconvolution (CSD) using the *tournier* algorithm (Tournier, Calamante, & Connelly, 2013); and (3) FOD estimated with the multi-shell multi-tissue CSD (msmt CSD) using the *dhollander* algorithm, which provides an unsupervised estimation of tissue specific response functions. The msmt CSD was performed using a WM/CSF compartment model (Jeurissen, Tournier, Dhollander, Connelly, & Sijbers, 2014). Each of the three tractography models was then paired with a deterministic and a probabilistic tracking algorithm. Deterministic DT-based tracking was performed using Euler integration (*Tensor_Det*; (Basser, Pajevic, Pierpaoli, Duda, & Aldroubi, 2000), while DT-based probabilistic tracking used bootstrapping (*Tensor_Prob*; (Jones, 2008). CSD-based tractography was performed according to FOD peaks either deterministically (SD_STREAM; (Tournier, Calamante, & Connelly, 2012) or probabilistically (iFOD2; (Tournier, Calamante, & Connelly, 2010). A spherical harmonic order of 8 was used for CSD-based estimations. One million streamlines were tracked over the full brain with the parameters recommended by MRtrix3: stepsize 0.024 mm (0.12 mm for iFOD2), angle 90° per voxel (45° for iFOD2), minimal streamline length 1.2 mm, maximal length 2.4 cm. Streamline seeds were produced at random locations within the brain mask until the defined number of streamlines was reached. To prevent streamlines from going across sulci, the brain mask was used as a stopping criterion.

### Tractography based connectivity matrices

Structural connectivity matrices were extracted from the tractography results using the number of streamlines connecting pairs of regions. The connectivity matrices are available in the supplementary material (Supplementary file 1). Matrices reporting the averaged fiber lengths between regions were also computed. Then, structural connectivity matrices were normalized using fractional scaling, such that the number of streamlines between pairs of regions were divided by the sum of the streamline counts connected to each of the regions, excluding self-connections (Donahue et al., 2016). The weights then represent the fraction of streamlines (FS).

All MRI data analysis was performed using the MRtrix3 software (http://www.mrtrix.org/), and custom scripts for Python (www.python.org), including python packages Nipype (Gorgolewski et al., 2011), Nibabel (Brett et al., 2018) and Numpy (Oliphant, 2015). All the scripts and data are available on the following GitHub repository: https://github.com/neuroanatomy/FerretDiffusionTractTracingComparison.

### Anatomical tract-tracing data

Structural connectivity data from anatomical tract-tracing experiments in adult ferrets (of two years age) were obtained from (Dell et al., 2019a, 2019b, 2019c). The experiments examined the cortico-cortical and cortico-thalamic connectivity of areas 17, 18, 19 and 21 (occipital visual cortex), PPc and PPr (posterior parietal visual cortex), and 20a and 20b (temporal visual cortex) in adult ferrets by means of retrograde Biotinylated Dextran Amine tracer (BDA). By retrograde tract-tracing, neuronal projections were traced from the axon terminations located in areas 17, 18, 19, 21, PPc, PPr, 20a and 20b (injection sites) of one hemisphere to the neurons’ cell bodies, located in different brain regions and across both brain hemispheres. Thus, the injected brain regions were defined as the target regions and the brain regions with cell bodies that were labeled positive for BDA were defined as the source regions. The connections were then quantified by obtaining a fraction of labelled neuron (FLN) value; refer to (Dell et al., 2019a, 2019b, 2019c) for a detailed explanation on the experimental procedures. Furthermore, for the purpose of this study, we considered only ipsilateral projections and adjusted the connectivity matrix and FLN values to exclude contralateral projections, in order to focus on the edge-complete subnetwork.

### Tract-tracing based connectivity matrix

A structural connectivity matrix was assembled from the left hemisphere such that the weights represent the number of retrograde labeled neurons between pairs of regions. This provided us with an asymmetric (directed) matrix indicating projections to the tracer injection sites. The weights were normalized using the fraction of labeled neurons (FLN), the number of labeled neurons in a source region divided by the total number of labeled neurons from the injected region (Markov et al., 2014). Considering that diffusion MRI tractography does not provide information on the directionality of the connections, the tract-tracing matrix was also symmetrized by averaging FLN values in both directions.

### Statistical analyses

Correlation coefficients were used to quantify the degree to which diffusion MRI tractography matched tract-tracing data. Thereafter, in order to characterize the ability of tractography to map structural weights, the strongest connections in the tract-tracing data were progressively removed from both sources (tractography and tract-tracing), and correlation coefficients were then computed on the remaining connections. In the same way, we also computed correlation coefficients when excluding the weakest tract-tracing connections. Such exploration allowed us to probe whether the correlation coefficient values were mainly driven by strong connections, which correlate with short-range connections and are statistically more likely to be detected. By contrast, weak or longer-range connections are more likely to be spurious (false positives). In order to deal with the log-normal distribution of structural connectivity values in both diffusion MRI tractography and tract-tracing experiments, we computed either the non-parametric Spearman’s correlation coefficient or the Pearson’s correlation coefficient on the values logarithmically transformed (both FLN and FS). In order to cope with absent connections when performing the logarithmic transformation, for the Pearson’s correlations, all raw counts of streamlines and labeled neurons (before the normalizations) were incremented by one. Confidence intervals were computed using bootstrapping at a confidence level of 95%. In addition, we computed the partial Spearman correlations when regressing out the euclidean distance between the centroids of our cortical areas. We first modelled the relationship between the logarithm of the FLN and FS values with the euclidean distance between each pair of cortical areas and extracted its residuals. The residuals from the FLN and the FS were then correlated using Spearman’s correlation.

To quantify the ability of tractography to correctly detect existing tract-tracing connections, we computed basic classification performance measures: sensitivity, specificity and precision. Sensitivity quantifies how good a measure is at detecting true connections, while specificity estimates how good a quantity is at avoiding false detections. Average precision quantifies how many of the positively detected connections were relevant. Tract-tracing structural connectivity matrix was progressively thresholded and binarized keeping a given proportion of the strongest weights, from 0.1 to 0.9 by step of 0.1 (Rubinov & Sporns, 2010) in order to build a series of receiver operating characteristic and precision and recall curves. The performance measures were then averaged for each threshold as summary statistics.

The statistical analyses were performed using R (https://www.R-project.org/) and Python with the scikit-learn package (Garreta & Moncecchi, 2013).

## Results

Structural connectivity estimates from diffusion MRI tractography were all highly positively correlated with the tract-tracing data (Spearman’s rho ranging from 0.67 to 0.91, all p < 10^−3^) (Table 1 and Figure 1). Probabilistic tractography algorithms increased the correlation values obtained with deterministic tractography. The DT model was not able to recover all the connections found in tract-tracing data for both deterministic (7 connections) and probabilistic (5 connections) tractography, as shown by the ‘white’ circles that correspond to connections that were not found by the diffusion MRI tractography (lowest rank), and hence the average streamline length of these connections is zero (white circles) (Figure 1C and D). The 95% confidence intervals for the relative predictive power of the different tractography algorithms overlapped, suggesting an absence of statistically significant differences. Consistent results were observed when using the Pearson correlation coefficient (Table 1 and Supplementary Figure 1).

**Table 1:** Correlations between diffusion MRI tractography and tract-tracing experiments. P-values inferior to 1.10^−3^ are indicated by ** and p-values inferior to 0.05 by *.

|  |  | Undirected tract-tracing matrix |  | Directed tract-tracing matrix |  |
| --- | --- | --- | --- | --- | --- |
|  |  | Spearman | Pearson | Spearman | Pearson |
| Deterministic | DTI | 0.67 ** [0.44-0.94] | 0.69 ** [0.37-0.86] | 0.50 * [0.22-0.82] | 0.48 * [0.07-0.75] |
|  | CSD | 0.76 ** [0.56-1.00] | 0.68 ** [0.36-0.86] | 0.62 * [0.35-0.93] | 0.53 * [0.13-0.78] |
|  | msmt CSD | 0.71 ** [0.49-0.99] | 0.71 ** [0.40-0.87] | 0.57 * [0.22-0.95] | 0.55 ** [0.16-0.79] |
| Probabilistic | DTI | 0.79 ** [0.65-0.98] | 0.78 ** [0.53-0.90] | 0.67 ** [0.49-0.91] | 0.63 ** [0.27-0.83] |
|  | CSD | 0.91 ** [0.82-1.00] | 0.88 ** [0.73-0.95] | 0.77 ** [0.56-1.00] | 0.69 ** [0.38-0.86] |
|  | msmt CSD | 0.87 ** [0.75-1.00] | 0.89 ** [0.76-0.95] | 0.70 ** [0.46-0.99] | 0.67 ** [0.33-0.85] |

Spearman correlations were decreased after regressing out the euclidean distance. Partial Spearman correlation values were no longer statistically significant for deterministic tractography (DTI: r = 0.36, p = 0.10; CSD: r = 0.39, p = 0.09; msmt CSD: r = 0.40, p = 0.07). However, for probabilistic tractography correlations remained statistically significant (DTI: r = 0.54, p < 0.05; CSD: r = 0.66, p < 0.05; msmt CSD: r = 0.77, p < 0.05), see supplementary figure 8. Consistent results were observed when using the Pearson correlation coefficient (Supplementary table 1).

We then tested the influence of strong and weak connections on the relationship between diffusion MRI tractography and tract-tracing data. Structural connectivity estimates from diffusion MRI tractography remained highly positively correlated to tract-tracing data after progressive removal of 25% of the strongest connections and similarly after removal of the weakest connections (Figure 2 and Supplementary Figure 2 for Pearson correlations). These results show that the correlations between diffusion tractography and tract-tracing were not primarily driven by connections most likely to be recovered by diffusion tractography because of their topographic proximity or their strength (strong weights). Similarly, we observed that the correlations were not affected by the weakest connections which are generally more sensitive to noise (leading to false positives), otherwise there would have been an increase in correlation values.

**Figure 2:**
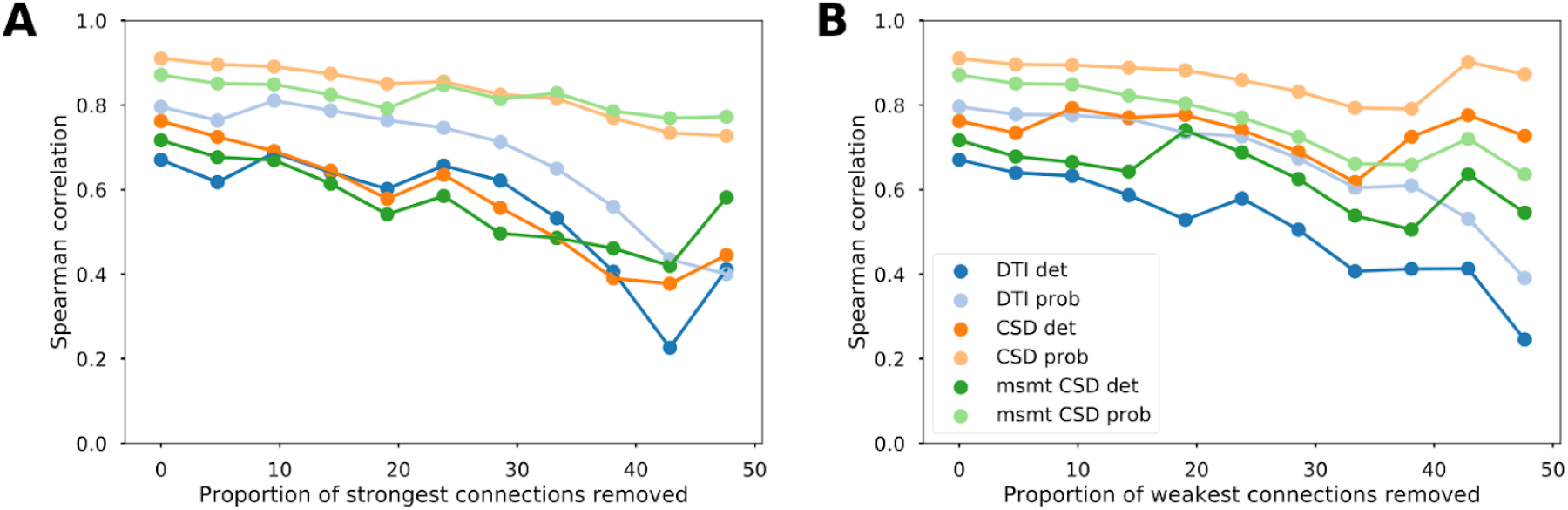
Reliability of the association between diffusion MRI tractography and tract-tracing data. Evolution of the Spearman correlation values between tract-tracing and diffusion MRI tractography data as a function of the proportion of strongest (A) and weakest (B) connections removed for the different tractography algorithms.

Measures of sensitivity/specificity/precision give an indication of the detectability of the connections. Our results were averaged and plotted as a function of the proportion of tract-tracing connections (Figure 3). CSD-based algorithms had generally higher sensitivity and precision compared to the diffusion tensor model, while tensor-based tractography had slightly higher specificity.

**Figure 3:**
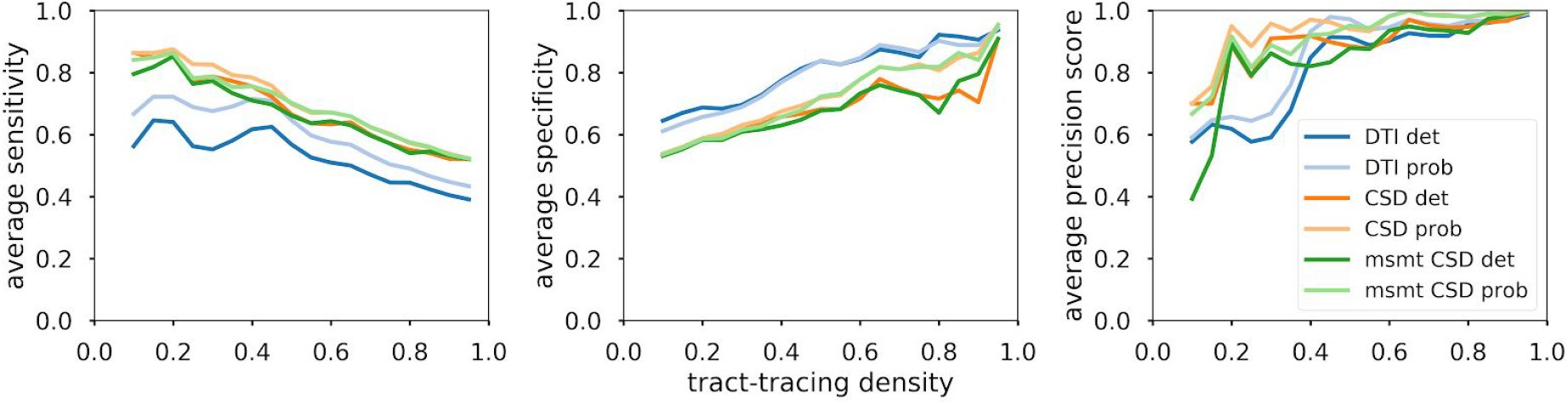
Detection performance of diffusion MRI tractography algorithms. Averaged sensitivity (A), specificity (B) and precision (C) as a function of the tract-tracing density.

All analyses were also performed comparing tractography with the directed structural connectivity from tract-tracing. We found decreased yet still statistically significant associations (Supplementary figures 3 to 7).

## Discussion

In the present study, we investigated the ability of different diffusion MRI tractography algorithms to reliably map ferret brain structural connectivity as retrieved from invasive tract-tracing experiments. We found that structural connectivity estimates from tractography were highly correlated with tract-tracing data. The different algorithms presented small, non-significant variations. Our results confirm in the ferret results from previous studies in the monkey (Donahue et al., 2016) and the mouse (Calabrese et al., 2015) as well as results using manganese tracing in the pig (Knösche et al., 2015). Overall, these findings enhance our confidence in diffusion MRI tractography as a powerful tool for exploring the structural connectional architecture of the brain.

We obtained estimates of the reliability of six different tractography algorithms with regard to tract-tracing data for the same cortical areas of the ferret brain. CSD-based algorithms presented the highest degree of concordance with tract-tracing data, and DT-based algorithms the least. However, the differences in correlation values did not appear to be statistically significant, as suggested by the overlapping 95% confidence intervals. High concordance with no particular algorithm outperforming the others was also reported when matching tract pathways from tractography and manganese tracing for a set of tractography algorithms (Knösche et al., 2015). Comparable overall correlations of the weighted connections have been obtained in the macaque brain, with a Spearman’s correlation of 0.59 (Donahue et al., 2016). However, here we report little effect of the strongest/weakest connections in the correlation values. The gradual decrease of our correlations indicated that our correlations were not amplified by the weight of strong connections or underestimated by a high amount of false positives stemming from weak connections. In addition, we showed high detection performance values across algorithms. Consistent with the correlation analysis, we observed higher performances for CSD-based algorithms in terms of precision. Also consistent with prior studies, DT-based results appeared to give slightly higher specificity than CSD-based algorithms, to the detriment of its sensitivity (Knösche et al., 2015; Sarwar et al., 2018). Such results are likely due to the lower ability of diffusion tensor models to resolve complex fiber geometries (Maier-Hein et al., 2017; Zalesky et al., 2016).

Our correlations were decreased and no longer statistically significant after regressing out distance, for deterministic tractography. Similar results have been reported in the macaque, where correlations decreased from r = 0.59 to r = 0.22 after regressing the distance effects (Donahue et al., 2016). Tractography’s ability to recover tracts is expected to decrease as a function of the distance due to technical biases (eg., in probabilistic tractography, the probability to follow a given path drops exponentially with distance). Thus, it has been shown that structural connectivity estimates from diffusion MRI tractography are highly related to their lengths (Liptrot, Sidaros, & Dyrby, 2014; Roberts et al., 2016). On the other hand, distance is a biological principle for the preferential connection between two brain areas (Hilgetag et al. 2016). As such, it remains challenging to disentangle these two factors from tractography outputs. Our regions can also be considered as neighbors relative to the whole brain size as they are all located in the occipital, parietal and temporal lobes of one hemisphere. This proximity could have inflated our correlations benefiting from the ability of tratography and tract-tracing to more accurately recover connections from neighboring areas. In any case, the correlations in which distance was regressed out, which corrects for both effects of distance (proximity and remoteness), maintain statistically significant correlations for all probabilistic tractography algorithms (especially based on CSD).

Our results showed a high correlation between diffusion MRI tractography and tract-tracing data, however, we note the limitations in our methodology. Firstly, the two datasets had different origins (i.e. the tract-tracing and tractography were not performed in the same animal) and the sample sizes were very small. Although the ferrets could all be considered mature in terms of brain development (Jackson, Peduzzi, & Hickey, 1989; Neal et al., 2007), the ferret used for the MR imaging was only two months old, while the animals used in tract-tracing were around 2 years old. This may have increased inter-individual variability and induced a bias in our cortical parcellations: although the sulcal and gyral patterns (used for cortical parcellation of MRI data, in relation to (Bizley & King, 2009) are unchanged after postnatal week 4, the ferret brain is still undergoing maturation and growth in all brain structures. The ferret brain growth reaches a plateau at postnatal week 24, however, the differences due to age should be only minor because the cortical architecture at two months of age resembles that at adult age (Jackson et al., 1989; Neal et al., 2007). Similarly, the cortex continues to undergo rostrocaudal expansion until postnatal week 24, after which the ferret brain reaches its adult size; however, previous studies have showed no significant changes of MRI-measured indices (Barnette et al., 2009; Neal et al., 2007). Although the brain of a two month old ferret is structurally similar to that of an adult brain, it still undergoes functional differentiation and pruning of connections, which could result in a minor shift in the placement of our cortical cytoarchitectonic parcellations and such parcellations can only be observed in histological sections but not in MRI scans.

Secondly, tract-tracing experiments, despite considered as ground-truth, are not exempted of limitations, such as the creation of false positive and false negatives, specificity of tracer and antibody used, spillage of tracer and passive diffusion (Heimer & Robards, 2013; Köbbert et al., 2000; Zaborszky, Wouterlood, & Lanciego, 2006). In addition, in this study we only considered the retrograde connections which are easier to quantify and neglected anterograde tracing results.

In sum, this study allowed us to validate structural connectivity estimates from diffusion MRI tractography by comparison with tract-tracing data in the ferret brain and it provided an estimation of the performances of three diffusion tractography algorithms, namely DT, CSD and msmt CSD, using both deterministic and probabilistic tracking. Generally, the currently available connectivity data for the ferret is quite limited; therefore, whole-brain tractography based on diffusion imaging can provide an initial, worthwhile estimate of structural connectivity that can be used for further anatomical, developmental and computational studies of the ferret brain.

## Acknowledgements

We gratefully acknowledge financial support by grants from the Deutsche Forschungsgemeinschaft (DFG): SFB 936/A1/Z3 and SPP 2041 / HI 1286/6-1, the Human Brain Project HBP-SGA2 (785907)/ SGA2 as well as 2015 FLAG-ERA Joint Transnational Call for project FIIND, in particular, ANR-15-HBPR-0005.

