## Supplementary figures for "Comparison between diffusion MRI tractography and histological tract-tracing of cortico-cortical structural connectivity in the ferret brain"

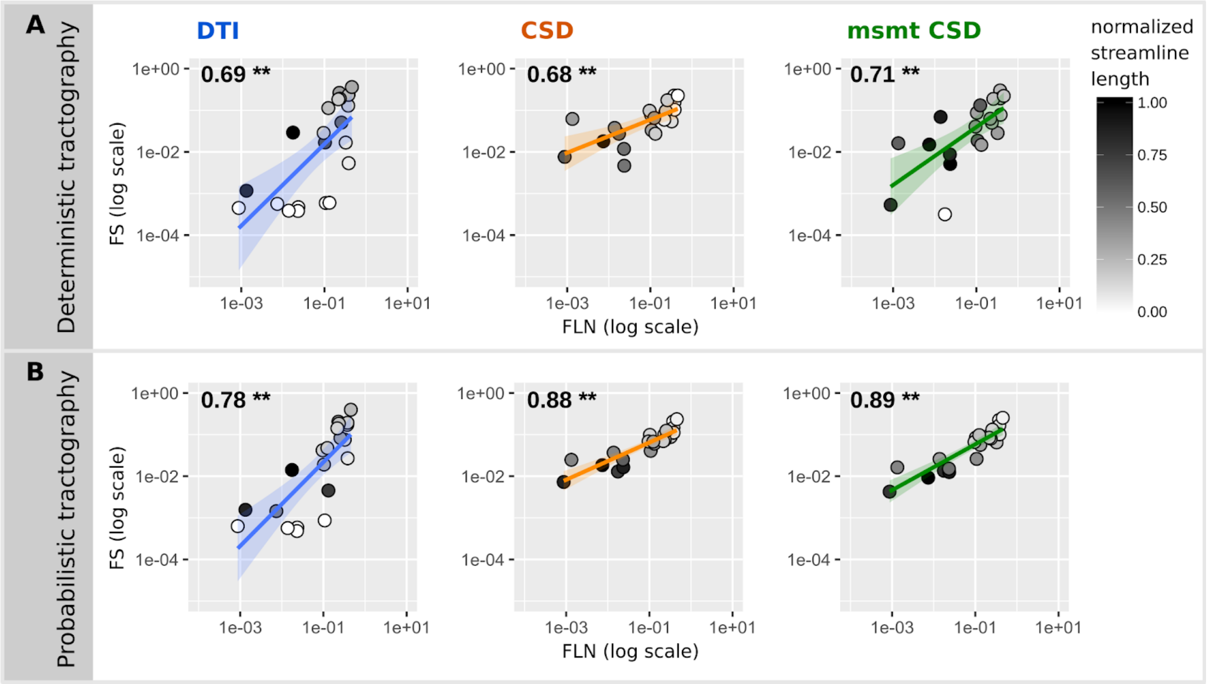


**Supplementary figure 1: Pearson's correlation between diffusion MRI tractography and tract-tracing experiments (symmetrical matrix).** Scatterplots of the ranked fraction of neurons vs. the ranked fraction of streamlines for deterministic (A) and the probabilistic (B) tractography. Grey colors code for the average streamline length (values normalized by the maximum streamline length of all the algorithms). P-values inferior to $1.10^{-3}$ are indicated by ** and p-values inferior to 0.05 by *.

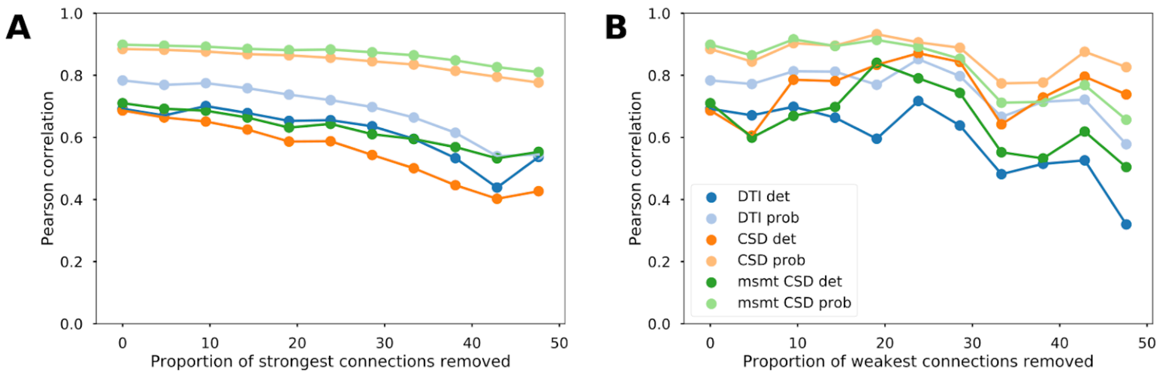


**Supplementary figure 2: Reliability of the association between diffusion MRI tractography and tract-tracing data (symmetrical matrix).** Evolution of the Pearson correlation values between tract-tracing and diffusion MRI tractography data as a function of the proportion of removed strong (A) and weak (B) connections for the different tractography models.

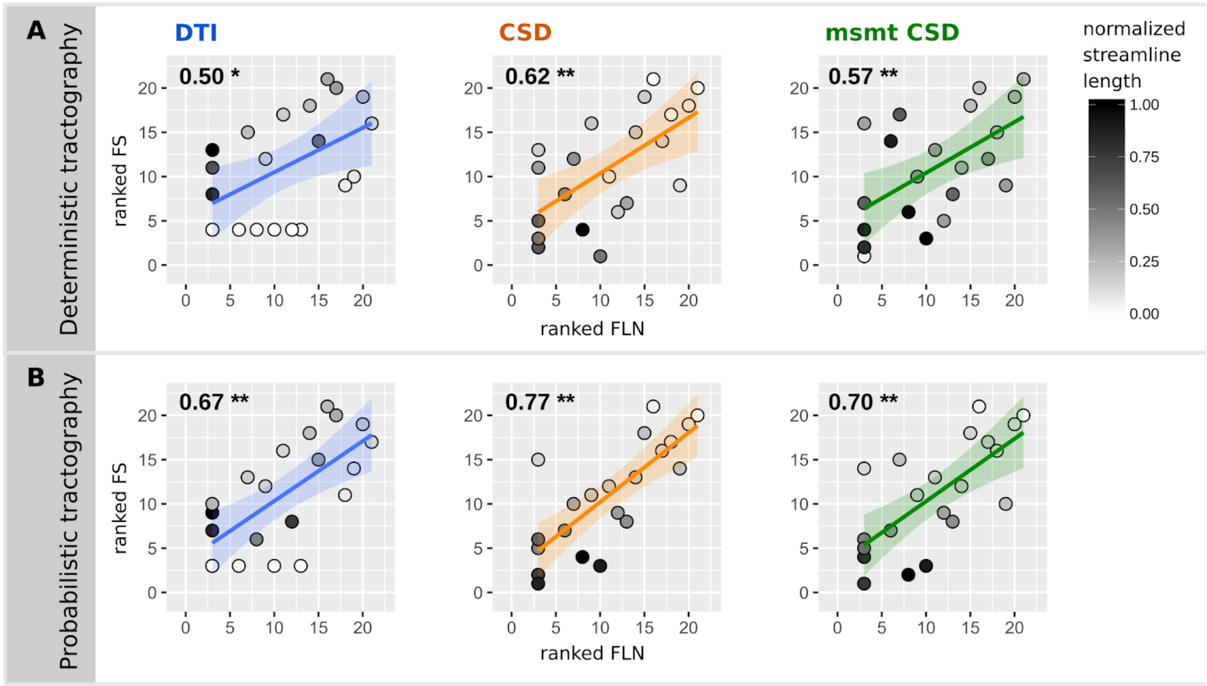


**Supplementary figure 3: Spearman's correlation between diffusion MRI tractography and tract-tracing experiments (directed matrix).** Scatterplots of the ranked fraction of neurons vs. the ranked fraction of streamlines for deterministic (A) and the probabilistic (B) tractography. Grey colors code for the average streamline length (values normalized by the maximum streamline length of all the algorithms). P-values inferior to $1.10^{-3}$ are indicated by ** and p-values inferior to 0.05 by *.

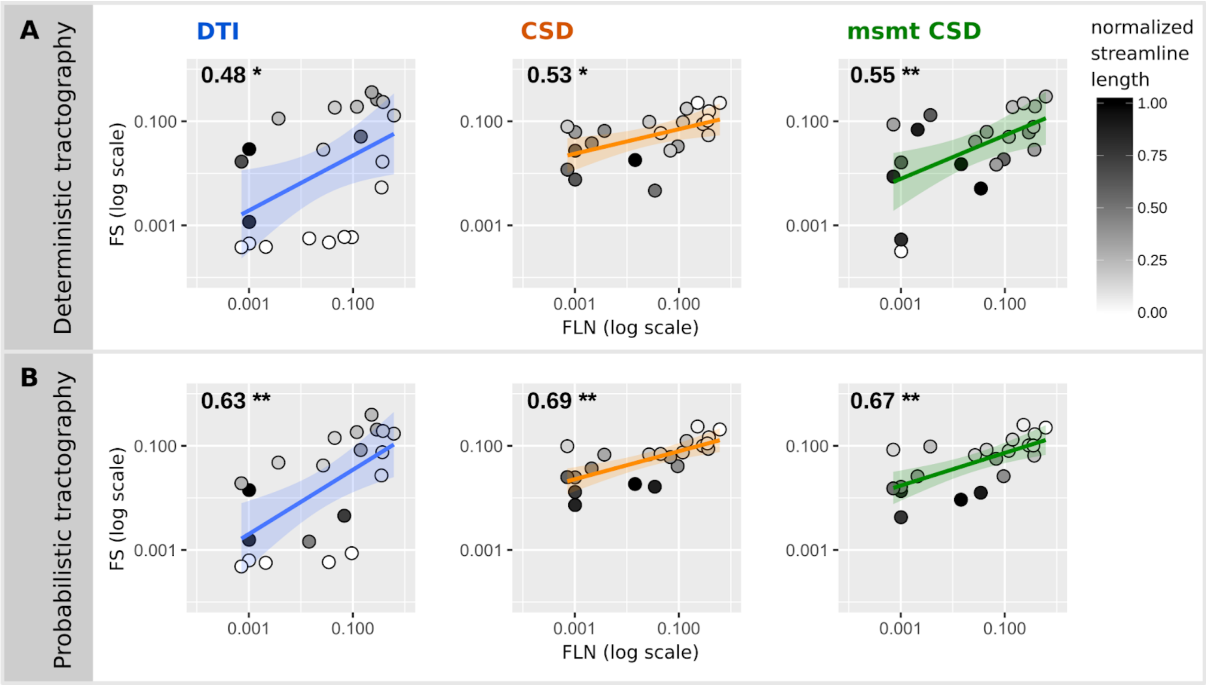


**Supplementary figure 4: Pearson's correlation between diffusion MRI tractography and tract-tracing experiments (directed matrix).** Scatterplots of the ranked fraction of neurons vs. the ranked fraction of streamlines for deterministic (A) and the probabilistic (B) tractography. Grey colors code for the average streamline length (values normalized by the maximum streamline length of all the algorithms). P-values inferior to $1.10^{-3}$ are indicated by ** and p-values inferior to 0.05 by *.

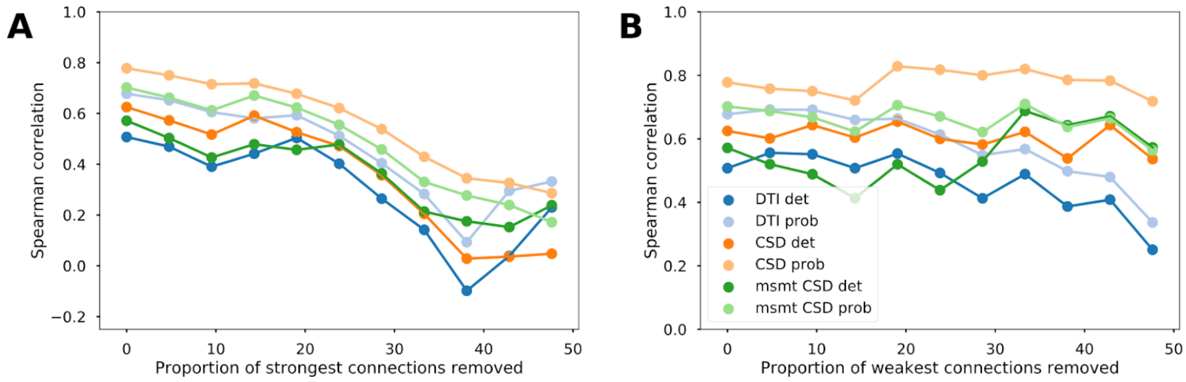


**Supplementary figure 5: Reliability of the association between diffusion MRI tractography and tract-tracing data (directed matrix).** Evolution of the Spearman correlation values between tract-tracing and diffusion MRI tractography data as a function of the proportion of removed strong (A) and weak (B) connections for the different tractography models.

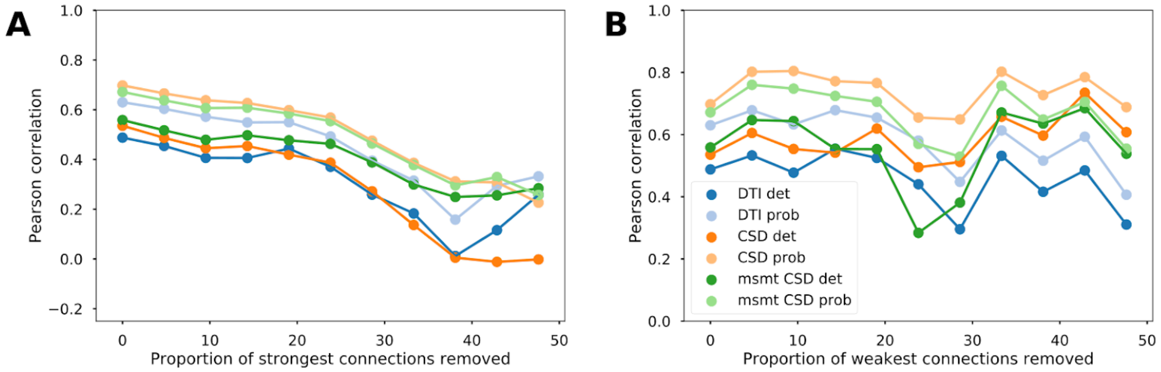


**Supplementary figure 6: Reliability of the association between diffusion MRI tractography and tract-tracing data (directed matrix).** Evolution of the Pearson correlation values between tract-tracing and diffusion MRI tractography data as a function of the proportion of removed strong (A) and weak (B) connections for the different tractography models.

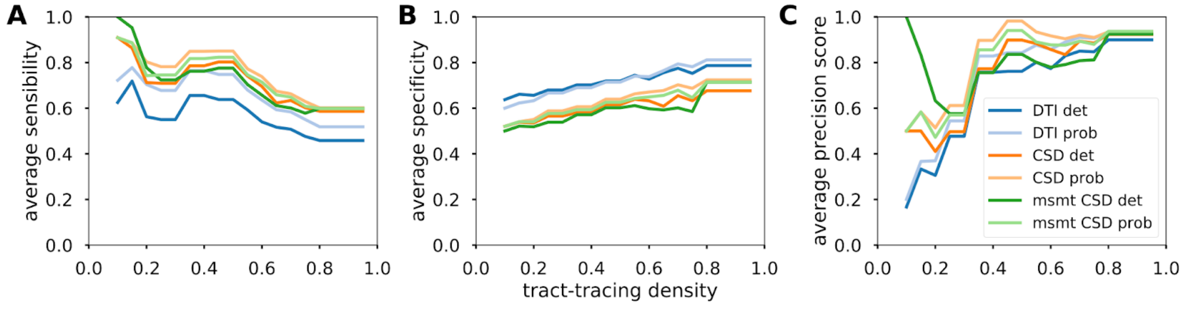

**Supplementary figure 7:** Average sensitivity (A), average specificity (B) and average precision score (C) along tract-tracing density (directed matrix).

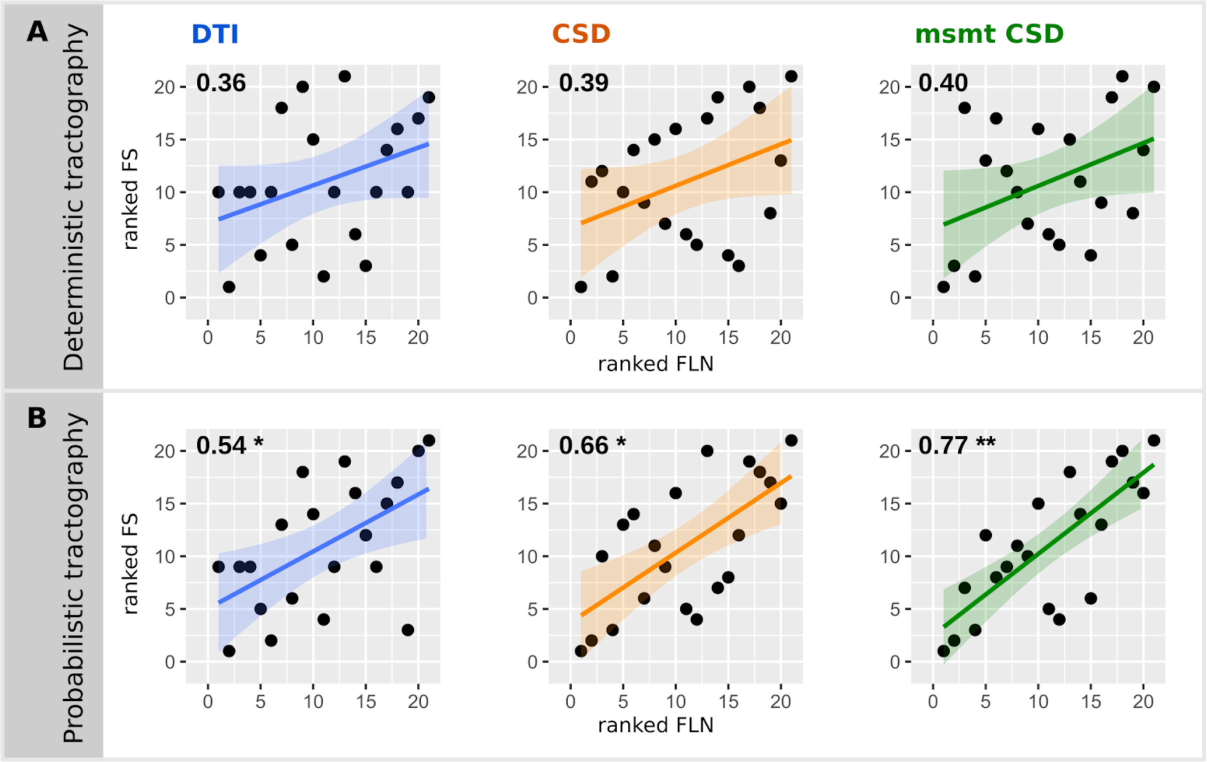


**Supplementary figure 8: Spearman's partial correlation between diffusion MRI tractography and tract-tracing experiments (symmetrical matrix).** Scatterplots of the ranked residuals FLN after regressing out the euclidean distance between each pair of areas vs. the ranked residuals FS for deterministic (A) and the probabilistic (B) tractography. P-values inferior to $1.10^{-3}$ are indicated by ** and p-values inferior to 0.05 by *.

| | | Undirected tract-tracing matrix | | Directed tract-tracing matrix | |
|---|---|---|---|---|---|
| | | Spearman | Pearson | Spearman | Pearson |
| Deterministic | DTI | 0.36 | 0.56 * | 0.38 | 0.3 |
| | CSD | 0.39 | 0.44 * | 0.35 | 0.23 |
| | msmt CSD | 0.4 | 0.50 * | 0.39 | 0.3 |
| Probabilistic | DTI | 0.54 * | 0.66 ** | 0.55 * | 0.46 * |
| | CSD | 0.66 * | 0.85 ** | 0.67 * | 0.53 * |
| | msmt CSD | 0.77 ** | 0.85 ** | 0.77 ** | 0.48 * |

**Supplementary table 1: Partial correlations between diffusion MRI tractography and tract-tracing experiments after regressing out the euclidean distance between each pair of areas.**

**Supplementary file 1: Number of connections between each pair of areas recovered by diffusion MRI tractography for DTI, CSD and msmt CSD tracked with deterministic and probabilistic tractography.**
